## Supplementary figure 1 and 2 for "Enhancing Prosthetic Vision by Upgrade of a Subretinal Photovoltaic Implant in situ"

**Supplementary Figure. 1.** Toluidine blue stained histological section 6 weeks post-explantation. A thick acellular layer (yellow arrow) develops in the subretinal space (red dotted line marks the RPE/choroid boundary). The subretinal fibrosis is localized to the area where the implant was (yellow dotted lines mark the location of primary implant).

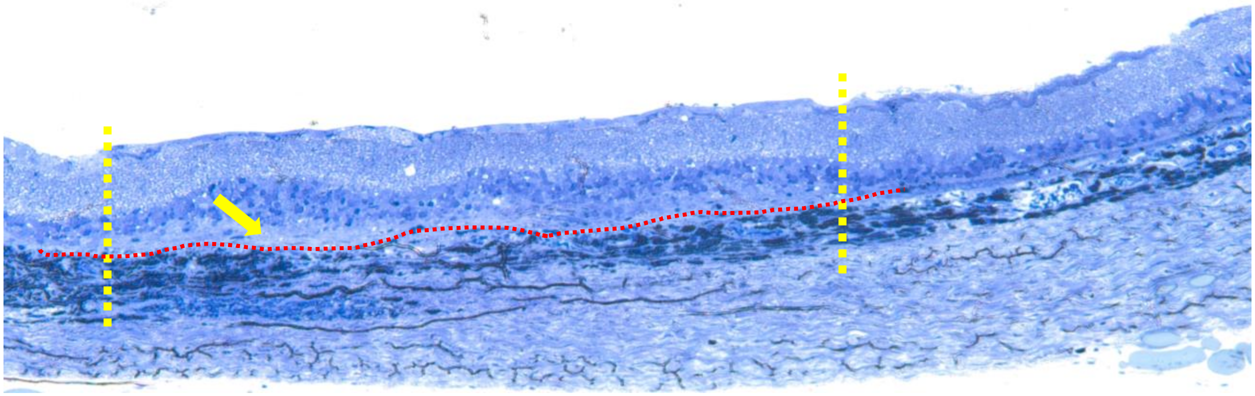

**Supplementary Figure. 2.** OCT image of an RCS control retina before surgery. The lens occupies the majority of the anterior chamber, leaving approximately 1 mm of space for retinal detachment, tool insertion and subretinal manipulations.

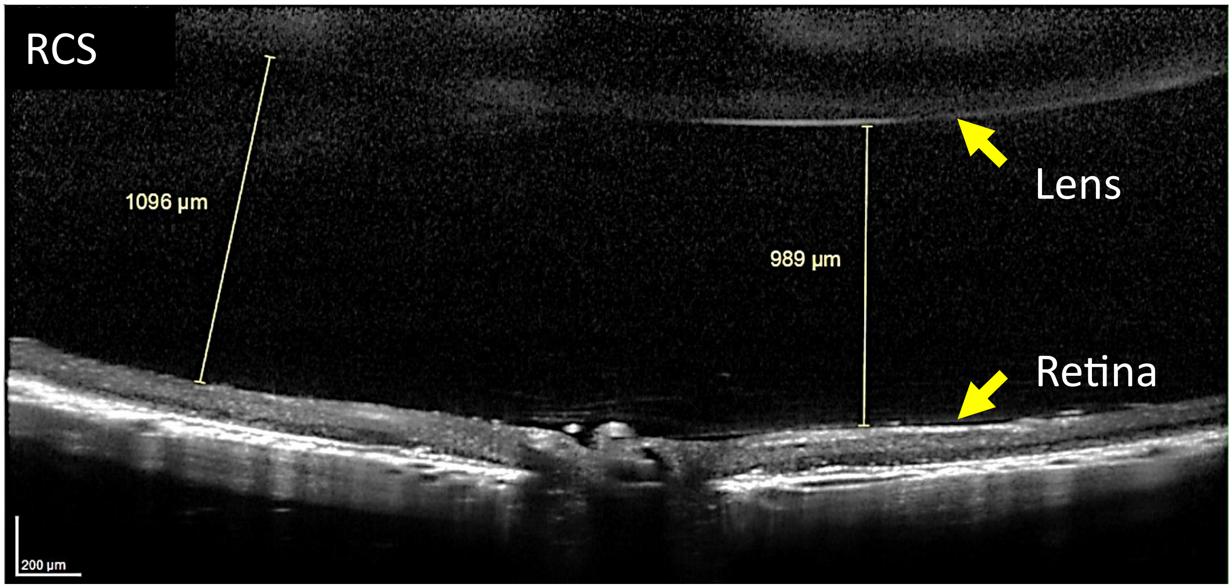
